# Comparative analysis of T-cell signatures and astroglial reactivity in Parkinson’s pathology across animal models with distinct regenerative capacities

**DOI:** 10.1101/2025.10.07.680901

**Authors:** Simona Intonti, Volker Enzmann, Amalia Perna, Ferdinando Spagnolo, Claudia Curcio, Federica Maria Conedera

## Abstract

Parkinson’s disease (PD) is a progressive neurodegenerative disorder characterized by the selective loss of dopaminergic (DAergic) neurons in the substantia nigra (SN) and the accumulation of misfolded α-synuclein (aSyn). Emerging evidence suggests that the immune system, particularly T-cell-mediated responses, plays a key role in the pathogenesis of PD. However, the heterogeneity of these immune responses across species and preclinical models with varying regenerative capacities remains poorly understood. In this study, we performed a comparative analysis of T-cell infiltration, astroglial reactivity, and DAergic neuronal loss across multiple models and species. These included acute DAergic degeneration induced by 1-methyl-4-phenyl-1,2,3,6-tetrahydropyridine (MPTP), genetically modified mice with accumulation of aSyn (Thy1-aSyn L61 model), adult zebrafish exposed to MPTP-induced neurotoxicity and human post-mortem midbrain tissue obtained from PD patients. Zebrafish exhibited transient DAergic neurodegeneration followed by neuronal regeneration and a temporary CD4^+^ T-cell infiltration alongside a regulated astroglial response. In contrast, MPTP-treated mice showed a permanent neuronal loss, increased astrogliosis and CD8^+^ T-cell infiltration that was negatively correlated with neuronal survival. In contrast, L61 mice exhibited progressive aSyn accumulation with chronic astrogliosis and CD4^+^ T-cell infiltration not directly linked to neuronal loss. Unlike age-matched controls, the SN from PD brains displayed DAergic degeneration, aSyn aggregation, and elevated CD3^+^ T-cell infiltration, which correlated with neuronal loss and aSyn burden. These findings emphasise the species- and model-specific immune profiles underlying PD pathology. Our results reveal that CD4^+^ T-cells contribute to neuronal regeneration following injury in zebrafish. This process is absent in the MPTP and L61 mouse models, which are instead driven by CD8^+^ or CD4^+^, respectively. This work underscores the potential of targeted immunomodulation aimed at T cell - glial interactions to slow neurodegeneration and promote repair in PD.

## INTRODUCTION

Neurodegenerative diseases are a group of disorders that affect the brain and nervous system, typically characterized by the progressive loss and dysfunction of neurons (1). Among progressive neurodegenerative diseases, Parkinson’s disease (PD) is the second most common condition (2). As the human population ages, the prevalence of PD is expected to rise significantly, with an estimated 25.2 million PD patients worldwide by 2050 (3).

PD is characterized by the loss of dopaminergic (DAergic) neurons in the substantia nigra (SN) and the accumulation of misfolded α-synuclein (aSyn), leading to motor and non-motor deficits in affected individuals (4). While the pathogenesis of PD is multifactorial, mounting evidence highlights the critical involvement of the immune system, particularly T-cell-mediated responses, in modulating neuronal degeneration and disease progression (5, 6). However, the heterogeneity of T-cell responses across different animal models and species with varying regenerative capacities remains poorly understood (7). Recently, not only microglia but also astrocytes have emerged as key contributors to PD pathology. In zebrafish, astrocyte-like radial glial cells respond to injury by actively supporting neuronal regeneration, providing a strong rationale for examining astroglial responses in comparative studies of PD (8). Glial cells, however, have a dual role in the repair process. In regenerative species, such as zebrafish, they enhance neuronal survival and regrowth, whereas in mammals, including mice, they frequently drive glial scar formation, thereby restricting axonal regeneration (9).

To induce PD-like features, we inject 1-methyl-4-phenyl-1,2,3,6-tetrahydropyridine (MPTP) in zebrafish and mice. This toxin-based model has long been a cornerstone of PD research because of its high reproducibility and ability to recapitulate some key neuropathological signatures of the human disease, including selective DAergic neuron loss and neuroinflammation (10, 11). In zebrafish, MPTP-induced neuron loss was followed by activation of the brain and retinal regenerative programs, which is characteristic of this species. In contrast, the same toxin caused an acute and irreversible degeneration of DAergic neurons in mice, reflecting the limited regenerative potential of the mammalian brain (11–13).

While the MPTP model reproduces the DAergic neurodegeneration observed in PD, it does not capture the progressive aSyn proteinopathy that is a defining feature of the human disorder. To address this limitation, we also used 4-month-old Thy1-aSyn (L61) transgenic mice that overexpress human aSyn but do not yet exhibit DAergic degeneration. While aSyn overexpression leads to early protein accumulation and synaptic alterations, motor dysfunction and DAergic neuron loss typically emerge after 14 months of age (14). Indeed, as the mice age, they develop widespread aSyn aggregates, particularly in cortical and subcortical regions, along with neuroinflammatory responses and impaired dopamine neurotransmission (15). Unlike the acute and rapid neuronal loss observed in MPTP-treated animals, the L61 model recapitulates the slow, cumulative, and aSyn-driven pathology of human PD (16). Together, these models provide a complementary framework to investigate how immune responses, astroglial activation, and regenerative capacity vary according to the nature and timing of the underlying insult. Lastly, to relate these experimental findings to human disease, we also examined post-mortem PD brains, where aSyn pathology is accompanied by immune activation. Building on emerging evidence for adaptive immune mechanisms in PD (17), we hypothesize that T cell infiltration of the nigrostriatal system drives the neuroinflammatory milieu, contributing to disease progression. In particular, T cells may exacerbate neurodegeneration, and modulation of their activity offers an attractive target for therapeutic intervention aimed at slowing the progression of PD.

To address this, we conducted a comparative study that investigates T-cell infiltration, phenotype, and distribution across established mammalian and non-mammalian models of PD, as well as in human post-mortem brain tissue from PD subjects and age-matched controls. We systematically profiled T-cells in the SN of MPTP-treated mice and zebrafish to capture toxin-induced injury, as well as L61 transgenic mice that mimic chronic a-synucleinopathy. These experimental data were then integrated with observations from human PD specimens to elucidate model- and species-specific patterns of T-cell involvement and explore how their immune repertoire and regenerative capacity shape the heterogeneity of T-cell outcomes observed in PD. Ultimately, our cross-species approach provides insight into T-cell-driven disease mechanisms that could guide the development of targeted immunomodulatory strategies with translational potential for patients with PD.

## MATERIALS AND METHODS

### Bibliographic data mining

We queried PubMed programmatically using the NCBI Entrez E-utilities for four experimental contexts: zebrafish, mouse, human, and cross-species comparative models of PD.

Each query was executed within an adaptive year-window recursion that partitions the global time span from January 1, 1960, to December 31, 2025. All PMIDs were harvested in batches to guarantee completeness and server compliance, yielding 35,502 unique PMIDs. For every PMID, we retrieved full MEDLINE XML and normalised title, abstract, MeSH, keyword, journal, authors and publication year into a relational dataframe (pandas v2.2) **(Suppl. Table 1)**.

The structured neuro-immunological information was extracted from the complete abstract set using DeepSeek-r1 (7-B, instruction-tuned), executed locally via Ollama on a CUDA 12.4 GPU (Quadro T2000, 4 GB VRAM). A deterministic, few-shot prompt constrained the model to populate a nine-field template (context, species, GFAP-CD4/CD8 relations, Dopaminergic neuron loss, multivariate pattern, GFAP reliability as a T-cell marker). Responses were parsed with regex against pre-validated patterns; if ≥5 fields were “not mentioned” the abstract was automatically re-submitted with a stricter fallback prompt. The pipeline yielded 380 high-confidence structured records. Outputs underwent (i) automatic sanity checks (duplicate abstracts, illegal field values, missing keys) and (ii) manual review of a 10 % stratified random sample per species. Discrepancies were corrected or flagged, concordance with human curators. By combining rule-based post-processing, duplicate removal and critical human validation, we minimised recency biases intrinsic to LLMs, ensuring a robust evidence base for downstream statistical and clustering analyses. The code is provided in the supplementary material.

### Zebrafish Studies

Adult zebrafish of both sexes (AB strain) were kept at the animal facility of the Department of Anatomy, University of Bern. Only adult zebrafish (∼1 year old) were used in this study. They were maintained under standard conditions in tank water at 26.5 °C and raised on a 14/10 h light/dark cycle, as previously described (18, 19). Fish were fed dry food twice daily (GEMMA Micro 300; Westbrook, ME, USA) and TetraMin Tropical Flakes (Delphin-Amazonia AG, Münchenstein, Switzerland) once per day, as well as Artemia salina once per day. During experiments, animals were kept in tank water.

Adult zebrafish received intraperitoneal (IP) MPTP to induce degeneration of DA neurons (Sigma-Aldrich, St. Louis, MO, USA). MPTP was freshly dissolved in sterile saline to a final concentration of 10 mg/mL. The injection volume was adjusted according to body weight, with each fish receiving 200 µg MPTP per gram body weight in a total volume of 10 µL per fish (11, 19, 20).

Prior to injection, fish were anesthetized with 0.16 mg/mL ethyl 3-aminobenzoate methanesulfonate salt (Tricaine; Sigma-Aldrich, Buchs, Switzerland) dissolved in tank water until loss of the righting reflex. Anesthetized fish were placed ventral side up on a moist sponge for stabilization. Using a 30-gauge insulin syringe, the MPTP solution was injected into the intraperitoneal cavity at the ventral midline, just posterior to the pelvic girdle, at an angle of approximately 45° to the body axis (11, 20). Control animals received an equivalent volume of sterile saline.

Following injection, fish were transferred to a recovery tank containing fresh system water and monitored until full recovery from anesthesia.

All experimental procedures were approved by the governmental authorities of the Canton of Bern (BE 58/2025).

### Mouse Studies

Adult C57BL/6J mice (Charles River Germany, Sulzfeld, Germany), both male and female, aged 6– 9 weeks, were used in all experiments. Animals were housed under standard conditions in individually ventilated cages (IVCs) with a 12-hour light/dark cycle in a temperature-controlled animal facility at the Central Animal Facilities of the University of Bern. Mice had ad libitum access to standard laboratory chow and water.

To induce DA neurodegeneration, mice received IP injections of MPTP at a dose of 40 mg/kg body weight, dissolved in sterile saline and administered in a total volume of 10 mL/kg (21). Control animals received an equivalent volume of sterile saline. All experimental procedures were approved by the governmental authorities of the Canton of Bern (BE 28/2024).

Paraffin-embedded 5 μm sections from 4-month-old Thy1-aSyn (L61) transgenic mice, which overexpress human wild-type aSyn under the Thy1 promoter, were kindly provided by Prof. Thomas Montine (Stanford University, Stanford, CA, USA) for comparative histological analysis.

### Human studies

Tissue microarrays (TMAs) of selected midbrain areas were produced as previously described (22–24). Briefly, archived human midbrain paraffin-embedded tissue was stained with hematoxylin-eosin (H&E) to identify regions of the left and right substantia nigra pars compacta (SNc) suitable for TMA construction. Tissue cores (2 mm diameter) were punched from these regions and transferred to recipient blocks, each containing approximately 40 samples. The first three punches in each TMA consisted of non-neuronal tissues (kidney or liver) as controls. Age-matched and PD samples were placed in separate blocks and sorted by collection date. TMA blocks were sectioned at 2.5 µm. The first section was stained with H&E, and the second was stained with aSyn to verify correct annotation and exclude aSyn pathology in age-matched samples.

All procedures involving human tissue were approved by the ethics committee of the Canton of Bern (KEK 200/14).

### Histology and Cell Quantification

Morphological analyses were performed on H&E-stained sections of eyes and brains collected at 1-, 7-, and 14-day post-injection (dpi) from C57BL/6J mice and zebrafish.

Following fixation in 4% paraformaldehyde at 4°C overnight, tissues were dehydrated in a graded ethanol series, cleared in xylene, and embedded in paraffin. Serial frontal sections were cut at a thickness of 5 µm using a rotary microtome, mounted on glass slides, deparaffinized, and rehydrated through a series of descending alcohol concentrations to water. H&E staining (hematoxylin from Sigma, St. Louis, MO, USA, and eosin from Roth, Karlsruhe, Germany) was performed according to established protocols (25).

Quantification of cell nuclei was carried out by manual counting on H&E-stained sections using a light microscope. In both zebrafish and mouse eyes, a region of interest measuring 200 µm in width was defined in the central retina, and all clearly identifiable nuclei within this region were counted (25). For mouse brain sections, the SN was identified based on anatomical landmarks, and all nuclei within this area were counted (26). In zebrafish brain sections, the posterior tuberculum (PT), considered functionally analogous to the mammalian SN (27), was similarly identified and analyzed. For each model, three non-overlapping sections were selected for analysis, and the average number of nuclei per region was calculated.

### Immunofluorescence

Paraffin sections (5 μm) were utilized for immunofluorescence. Antigen retrieval was achieved by incubating the sections in Tris–EDTA (pH 9.0) or Citrate buffer (pH 6.0) with 0.05% Tween-20 for 4 minutes, followed by cooling at room temperature for a minimum of 30 minutes. Blocking was performed for 1 h in Tris-buffered saline (TBS; pH 7.6) containing 5% goat normal serum (DAKO, Agilent Technologies, Baar, Switzerland) and 1% bovine serum albumin (Sigma-Aldrich). The sections were incubated overnight at 4 °C with primary antibodies diluted in TBS. Primary antibodies used in this study were rabbit anti-tyrosine hydroxylase (TH, 1:1000; Millipore, Billerica, MA, USA) rabbit anti-aSyn (phospho S129; ab51253; 10 μg/mL; Abcam, Cambridge, UK), rabbit anti-glial fibrillary acidic protein (GFAP; OPA1-06100; 1:200; Thermo Scientific, Wilmington, DE, USA), mouse anti-CD3 antibody (ab237721, 1:2000; Abcam), mouse anti-CD4 antibody (MA5-15775, Invitrogen, Carlsbad, CA, USA) and rabbit anti-CD8 antibody (ab217344, 1:1000; Abcam).

Sections were then washed for 20 minutes and stained at room temperature with the respective secondary Antibodies (1:500; Alexa Fluor 488 or 594, Abcam, Cambridge, UK) diluted in TBS with 1% BSA for 1 hour at room temperature. Cell nuclei were counterstained using Vectashield mounting media with 4′,6-diamidino-2-phenylindole (DAPI; Vector Labs, Burlingame, CA, USA).

### Statistics

All quantifications were performed by two independent, blinded observers using ImageJ/Fiji software (28) to minimize bias. Data were assessed for normality prior to statistical analysis. When normality was confirmed, parametric tests were applied, including unpaired t-tests or one-way ANOVA with Tukey’s post hoc test for multiple comparisons. If normality was not met, non-parametric tests such as the Mann–Whitney U test were used, and results are reported as median values. For longitudinal data, linear mixed-effects models were employed. The specific statistical tests used are detailed in the corresponding figure legends and text. Statistical significance was defined as p ≤ 0.05. All analyses were conducted using GraphPad Prism 9.4.1 (GraphPad Software, Boston, MA, USA). Correlation analyses were performed using Spearman’s correlation matrices generated with the corrplot function from the gplots package (version 3.0.1), and pairwise correlations between two variables were visualized using the ggscatter function from the ggpubr package (version 0.6.0), in R version 3.5.1.

## RESULTS

### Bibliographic data mining

Systematic parsing of 35 .502 Parkinson-related PubMed records yielded structured metadata for abstracts, authors and MeSH/keyword fields, exposing a species distribution dominated by human (26 .954), mouse (2. 558), comparative (5 .827) and zebrafish (163) studies and highlighting “Parkinson’s disease”, “α-synuclein”, “neuroinflammation” and “microglia” as the most frequent descriptors **(Fig. 1)**. Unsupervised text-mining (TF-IDF + PCA + K-means) resolved five stable thematic clusters and, through expanded term matching, quantified immune–degeneration markers across the corpus (GFAP = 1 .596, CD4 = 458, CD8 = 891, DA-loss = 6 .951 hits), while silhouette analysis (k = 5, score ≈ 0.998) confirmed cluster separability. Qualitative clustering delineated species-specific “information clouds” that can guide targeted literature searches and thereby refine subsequent content analysis **(Figure 1)**. A neuro-immunology-specialised Deepseek OLLAMA LLM prompt then captured 380 representative abstracts, showing a mouse-centred evidence base (58.3%) and revealing that only 23.1% explicitly report multivariate GFAP–CD4/CD8–CD8-dopaminergic-loss patterns, just 14 papers examine GFAP’s reliability as an astroglial marker, and 154 link CD4/CD8 infiltration to DA-neuron loss; full methodological details, cluster composition and PMID lists are provided in the “Bibliography exploitation” section of the Supplementary Materials.

**Figure 1:**
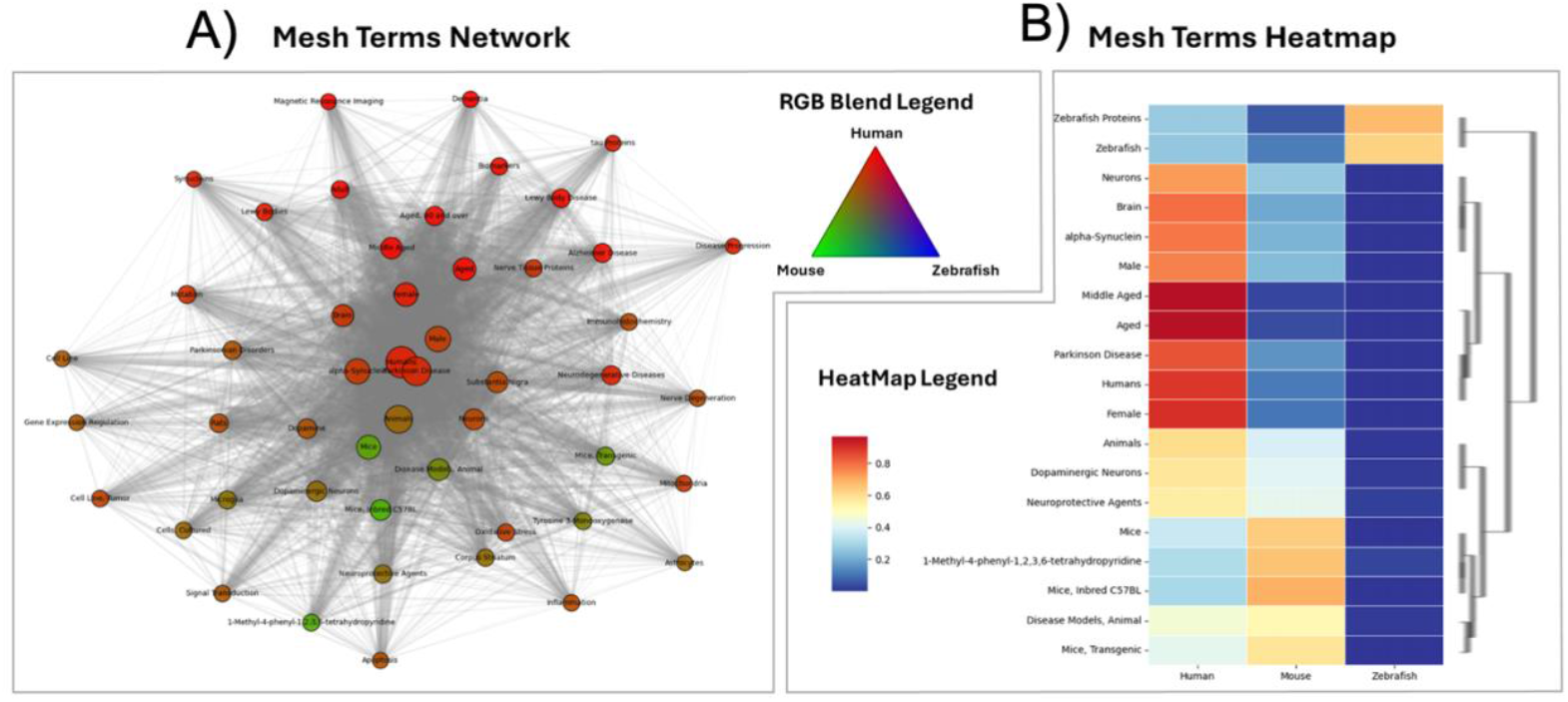
MeSH terms bibliographic relationships in Parkinson’s Disease models. Comparative analysis of the occurrences of MeSH terms associated with Parkinson’s Disease in the main experimental models (Human, Mouse, Zebrafish). **A)** The co-occurrence graph highlighting the relationships between the most representative terms (nodes RGB Blended according to their relative prevalence in the three species: red = Human, green = Mouse, blue = Zebrafish); **B)** Clustered heatmap with dendrogram showing the normalized distribution of the selected terms. Species-specific enrichment patterns can conceptually cluster clinical and neuropathological terms (Human), neuroinflammatory markers and transgenic models (Mouse), and regeneration and developmental terms (Zebrafish).

### Acute neurodegeneration in the Zebrafish model of PD

To reproduce PD-like neurodegeneration, adult zebrafish received IP injection of MPTP (200 μg/g), a dose previously shown to selectively damage DAergic neurons in zebrafish (11). We evaluated the effects of MPTP at 1, 7, and 14 days post-injection (dpi), focusing on two neuroanatomical regions highly sensitive to DA injury: the posterior tuberculum (PT), which is rich in DAergic neurons (29), and the retina. At 1 dpi, H&E staining of PT brain sections showed a non-significant decrease in the number of cells compared to the untreated control group **(Fig. 2A and B)**. At 7 dpi, the number of cells in the PT area significantly decreased compared to control (***p < 0.001), indicating a progressive neurodegeneration during the first week after MPTP injection **(Fig. 2A and B)**. Interestingly, the number of cells in the PT slightly increased at 14 dpi compared to 7 dpi, suggesting a partial recovery **(Fig. 2A and B)**. These results show that MPTP induces a transient neuronal damage in the zebrafish brain, followed by a regenerative process that becomes evident two weeks post-treatment. In parallel with brain neurodegeneration, the zebrafish retina exhibited layer-specific neuronal loss following MPTP exposure **(Fig. 2C and D)**. H&E staining showed a marked decrease in the number of cell nuclei in the inner nuclear layer (INL) at 1 and 7 dpi compared to the control group (**p < 0.01 and ***p < 0.001, respectively) **(Fig. 2D)**. At 14 dpi, cell numbers increased compared to 7 dpi. However, the difference was not statistically significant and reached control levels, suggesting a complete recovery of the INL. The ganglion cell layer (GC) and outer nuclear layer (ONL) remained unchanged, with no significant differences across all time points, suggesting that these retinal layers are less affected by MPTP treatment **(Fig. 2D)**. Overall, the data demonstrate that retinal neurons, particularly those in the INL, are highly vulnerable to MPTP-induced damage, but can undergo regeneration within two weeks post-treatment.

**Fig. 2:**
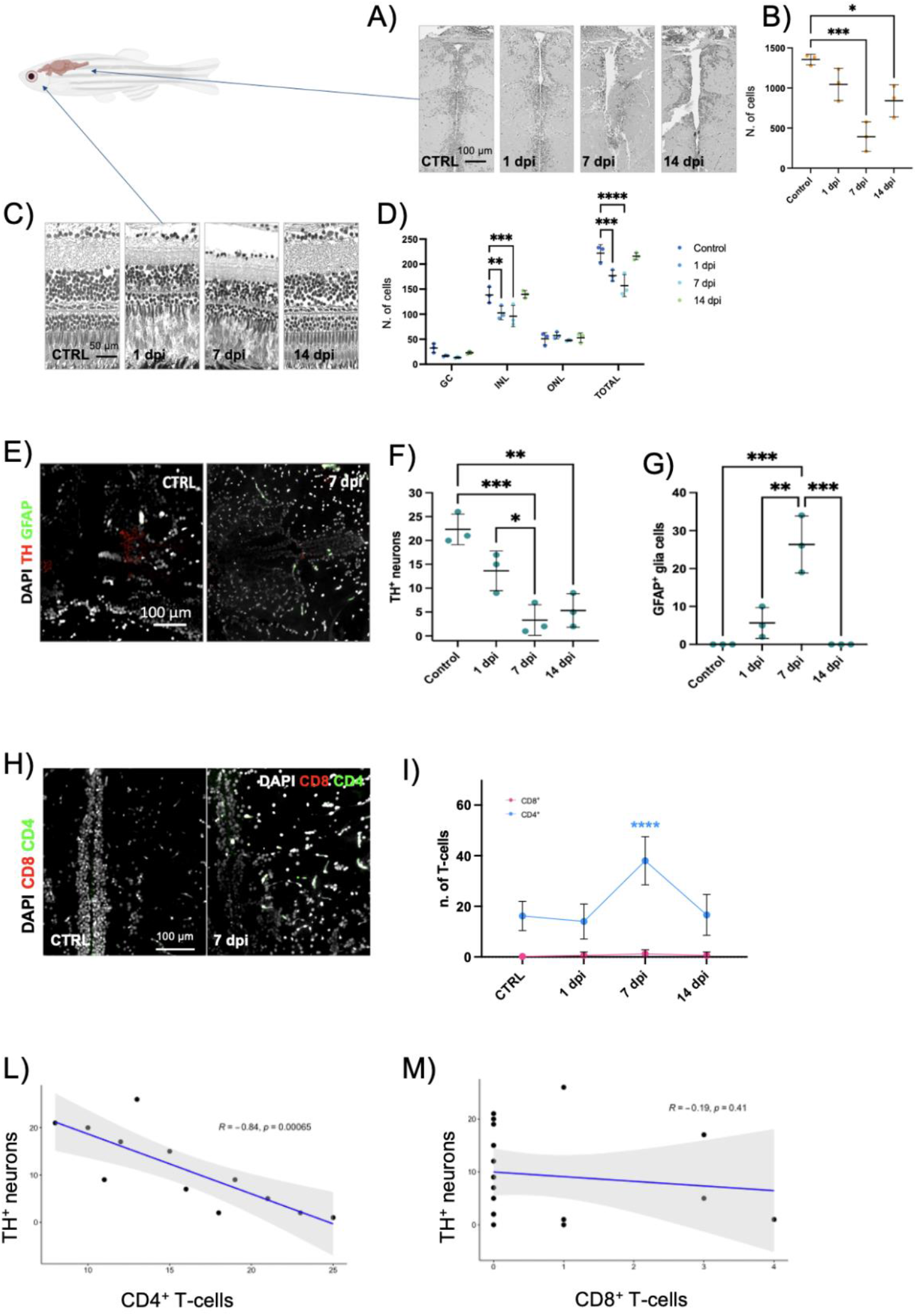
MPTP-induced neurodegeneration, astroglial activation, and T-cell infiltration in zebrafish. **A)** Representative H&E staining of zebrafish brain section showing PT in healthy control and MPTP-treated zebrafish at 1, 7, and 14 dpi; **B)** A reduction in the number of cells within the PT is observed at 7 and 14 dpi compared to controls, Quantification was performed using at least three animals per time point; **C)** Representative H&E-stained retinal sections from control and MPTP-treated zebrafish at 1, 7, and 14 dpi, highlighting structural alterations and cell loss across retinal layers over time; **D)** Quantification of retinal cell numbers shows a significant reduction at 1 and 7 dpi in the INL and total cell count. Quantification was performed using at least three animals per time point; **E)** Representative immunofluorescence images of zebrafish brain sections stained for DAergic neurons (TH, red) and glial (GFAP, green) cells at 1, 7, and 14 dpi with MPTP. Nuclei were stained with DAPI (white); **F)** Quantification of TH^+^ DAergic neurons and **G)** GFAP^+^ glial cells at 1, 7, and 14 dpi, Quantification was performed on at least three animals per time point, and statistical significance is shown in the graphs; **H)** Representative immunofluorescence images of the PT in control and MPTP-treated zebrafish at 7 dpi, stained for CD4^+^ (green) and CD8^+^ (red) T cells. Nuclei are stained with DAPI (white). **I)** Quantification of CD4^+^ and CD8^+^ T-cell subpopulations at different time points. The graph shows statistical significance. **L)** Statistical significance of the correlation between CD4^+^ and **M)** CD8^+^ T cells and a TH^+^ DAergic neurons is shown in the graphs. Statistical differences were assessed using one-way ANOVA and significance is shown in the graphs

To better characterize the involvement of astroglial response during MPTP-induced injury to DAergic neurons, we performed immunofluorescence staining using tyrosine hydroxylase (TH) to mark DAergic neurons (30) and GFAP as a marker for reactive astrocytes (31) **(Fig. 2E)**. At 7 and 14 dpi, the number of TH^+^ neurons were significantly lower than in control animals (***p < 0.001 and **p < 0.01 respectively) **(Fig. 2F)**. In parallel, a significant upregulation of GFAP^+^ cells at 7 dpi compared to controls was observed (***p < 0.001) **(Fig. 2G)**, suggesting a reactive astrogliosis peak. Interestingly, this glial response was followed by a significant decrease at 14 dpi compared to 7 dpi, reaching the baseline levels, indicating the resolution of glial reactivity **(Fig. 2G)**. Taken together, these results show that MPTP-induced DAergic neurodegeneration is accompanied by a transient glial activation, which peaks during the degenerative phase and as regeneration begins.

To evaluate the presence of T cell infiltration during MPTP-induced neurodegeneration, we performed an immunofluorescence staining using CD3, a canonical pan-T-cell marker, at different dpi, and we showed an increase of these cells at 7 dpi, followed by a decrease at 14 dpi, reaching the baseline level **(Suppl. Fig.1)**. To further characterize the T-cell heterogeneity we analyzed CD4^+^ and CD8^+^ subsets: Immunofluorescence analysis revealed an accumulation of CD4^+^ T cells at 7 dpi **(Fig.2H)**. In contrast, CD8^+^ T cells remained undetectable at every investigated timepoint **(Fig. 2H)**. Quantification confirmed a significant increase in CD4^+^ T cells at 7 dpi compared with control (****p < 0.0001), 1 dpi, and 14 dpi, suggesting a role for CD4^+^ T cells during the degenerative phase **(Fig. 1I)**. CD8^+^ T cells did not show any significant variation **(Fig. 2H and I)**. To investigate the relationship between different T cell subsets and neuronal damage, we performed a correlation analysis between CD4^+^ or CD8^+^ T cell counts and the number of DAergic neurons. A significant negative correlation was observed between CD4^+^ T cells and TH^+^ neurons (R = -0.84, p= 0.00065) **(Fig. 2L)**, while no significant correlation was found for CD8^+^ T cells **(Fig. 2M)**. These findings suggest that CD4^+^ T cells may play a role in promoting DAergic neuron loss or be involved in the subsequent regenerative process.

### Acute and chronic neurodegeneration in Mouse Models of PD

Adult wild-type mice were treated with MPTP using a protocol comparable to that used in zebrafish. We analyzed MPTP-induced neurodegeneration in the ventral midbrain region in mice, which corresponds to the posterior tuberculum (PT) in zebrafish, as well as the retina at 1-, 7-, and 14-days post-injection (dpi).

H&E staining of brain sections of MPTP-treated mice revealed a progressive loss of cellularity in the SN, initially observed at 1 dpi and further evident at 7 and 14 dpi **(Fig. 3A)**. Quantification of the number of cells showed a significant decrease at 1 dpi compared to controls (*p < 0.1) **(Fig. 3B)**. This reduction became more evident at 7 dpi and 14 dpi (***p < 0.001) **(Fig. 3B)**. These results confirm that MPTP-induced neurodegeneration in mice progresses over time without signs of spontaneous recovery. This contrasts with the regenerative response observed in zebrafish **(Fig. 2A and B)**, highlighting a species-specific difference in the ability to restore neural tissue following neurodegeneration induced by MPTP.

**Fig. 3:**
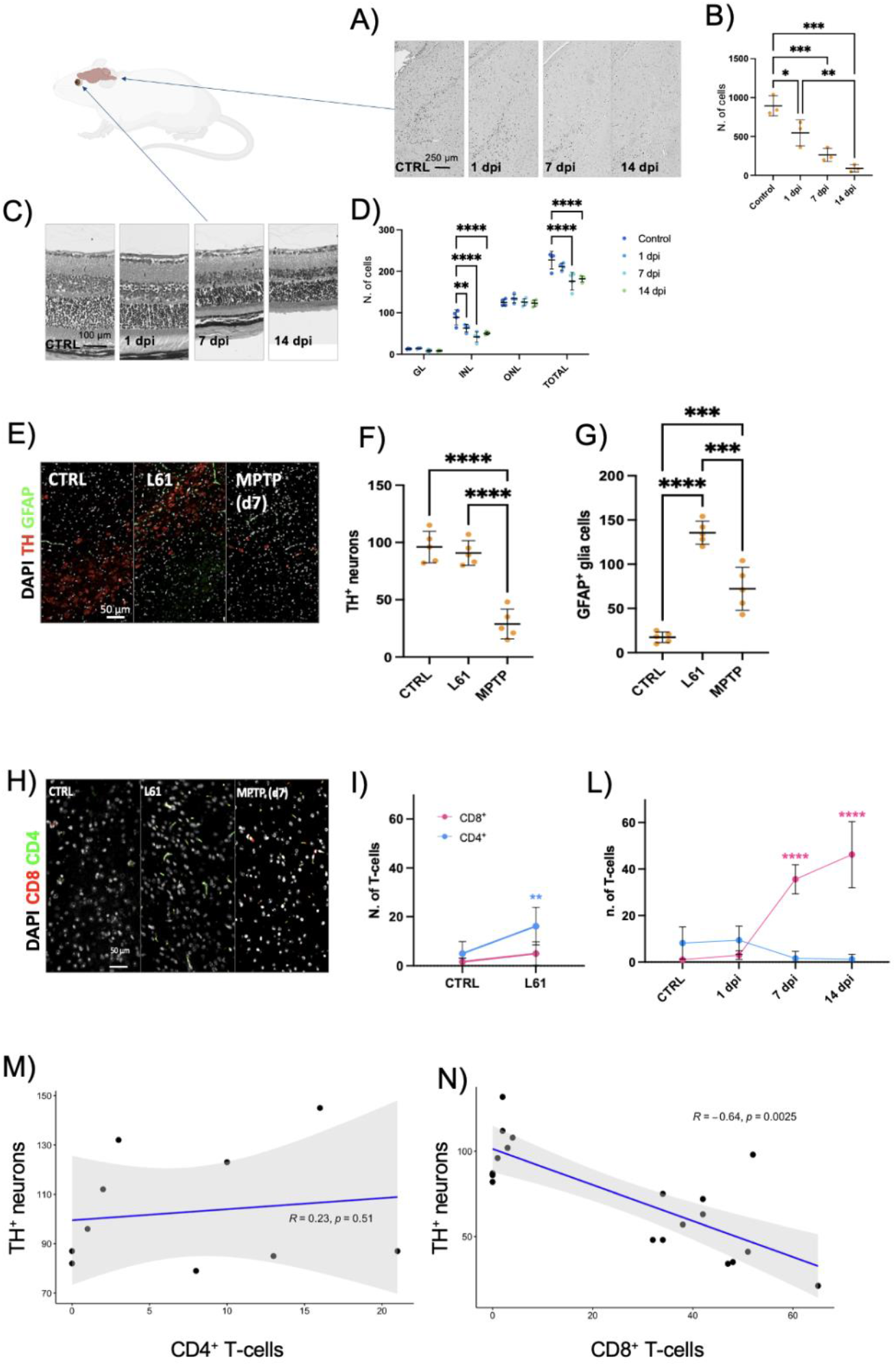
Neurodegeneration, Glial Activation, and T Cell Responses in Mouse Models of Parkinson’s Disease. **A)** Representative H&E-stained brain sections showing the SN in control and MPTP-treated mice at 1, 7, and 14 dpi; **B)** Quantification of the number of cells in the PT shows a gradual reduction over time following MPTP administration, with the lowest cell counts observed at 14 dpi. Statistical significance is shown in the graph; **C)** Representative H&E-stained retinal sections from control and MPTP-treated mice at 1, 7, and 14 dpi, showing progressive structural alterations and signs of recovery at 14 dpi; **D)** Quantification of cell numbers in the GL, INL, ONL, and total retina in MPTP-treated mice and controls. Quantification was performed on at least three animals per time point; **E)** Representative immunofluorescence images of SN from control, L61 (Thy1-aSyn) transgenic, and MPTP-treated mice. Sections were stained for DAergic neurons (TH^+^, red) and astroglial cells (GFAP^+^, green). Nuclei were stained with DAPI (white). Scale bar: 50 µm; **F)** Quantification of TH^+^ neurons and **G)** GFAP^+^ astrocytes in both L61 and MPTP-treated mice compared to controls; **H)** Representative immunofluorescence images of SN from control, L61, and MPTP-treated mice, stained for CD4^+^ (green) and CD8^+^ (red) T cells. Nuclei are stained with DAPI (white); **I)** L61 mice show predominantly CD4^+^ T-cell infiltration, while **L)** MPTP-treated mice exhibit a marked presence of CD8^+^ T cells; **M)** Correlation analysis between DAergic neuron number and CD4^**+**^ and **N)** CD8^**+**^ T cells in MPTP-treated mice. Statistical differences were assessed using one-way ANOVA, and significance is shown in the graphs

Retinal morphology was examined at 1, 7, and 14 days in wild-type mice following MPTP administration. Histological analysis with H&E staining **(Fig. 3C)** revealed evident tissue alterations that progressed over time. At 1 dpi, a reduction in cell density was evident, particularly affecting the INL. This loss became more pronounced by 7 dpi, with a marked thinning of the INL and disruption of the retinal structure. At 14 dpi, the retina looked similar to 7 dpi with no clear signs of recovery or further damage. In contrast, GL and ONL cell numbers remained like the control at all time points. To compare DAergic degeneration and astroglial reactivity elicited by two distinct PD-related insults, we analyzed the SN of L61 transgenic mice and MPTP-treated mice. Immunofluorescence analysis revealed a significant loss of TH^+^ DAergic neurons in 7 dpi MPTP-treated mice, whereas 4-month-old L61 transgenic mice showed no difference in the number of TH^+^ neurons compared to controls. Only the MPTP treatment caused a reduction in TH^+^ neuron numbers **(Fig. 3E, 3F)**. In parallel, we assessed astroglial activation using GFAP immunostaining **(Fig. 3G)**. Both models showed a significant increase in GFAP^+^ expression compared to the control, indicative of reactive astrogliosis. However, the astroglial response was more evident in L61 mice **(Fig. 3G)**, supporting that aSyn accumulation in this genetic model induces a stronger glial response than the acute MPTP-induced injury.

As in the zebrafish, we evaluated the presence of CD3^+^ T-cell infiltration in MPTP-treated and L61 mice by immunofluorescence and found that CD3^+^ T-cells were reduced at all time points analyzed **(Suppl. Fig. 2)**.

To further define the T-cell subtypes that infiltrate the SN during neurodegeneration, we analyzed CD4^+^ and CD8^+^ T-cells in L61 and MPTP-treated mice by immunofluorescence. Results revealed a difference in T-cell subtype distribution between the two mouse models, indicating that the nature and the stage of the neurodegenerative process affect the heterogeneity of T-cell infiltration in distinct ways. In L61 mice, the majority of infiltrating T cells were CD4^+^ (**p < 0.01), while CD8^+^ cells were absent **(Fig. 3H and I)**. Conversely, MPTP-treated mice showed a higher number of CD8^+^ T cells, especially at 7 and 14 dpi (**** p <0.0001), with only a few CD4^+^ cells observed **(Fig. 3H and L)**. To assess the potential contribution of T-cell subtypes to DAergic neurodegeneration in the MPTP model, we performed a correlation analysis between the number of infiltrating CD4^+^ and CD8^+^ T cells and the number of TH^+^ neurons in the SN of MPTP-treated mice. The analysis revealed a non-significant positive correlation between the number of CD4^+^ T cells and DAergic neurons **(Fig. 3M)**. In contrast, a significant negative correlation was observed between the number of CD8^+^ T cells and DAergic neurons (R = –0.64, p = 0.0025) **(Fig. 3N)**. This result indicates that higher infiltration of CD8^+^ T cells is associated with a DAergic neuron loss, supporting a potential cytotoxic role for CD8^+^ T cells in mediating neuronal damage.

### T cell infiltration in the SN of PD patients

To investigate immune involvement in PD, we examined SN sections prepared from post-mortem mesencephalon of PD patients and age-matched controls. We first assessed the extent of DAergic neurodegeneration by immunofluorescence staining on PD brains and observed a reduction and a disorganized distribution of TH^+^ neurons, compared with controls (**Fig. 4A**). In age-matched control sections, TH^+^ neurons appeared numerous, well organized, and well distributed throughout the SN (**Fig. 4A**). Quantitative analysis confirmed a significant decrease in TH^+^ cell numbers in PD tissues **(Fig. 4B)** compared to the control (****p <0.0001). Next, we assessed aggregated aSyn by immunohistochemistry for alpha-synuclein at serine 129 (pSer129), a pathological modification commonly associated with Lewy body formation and enriched in aggregated species. This accumulation is implicated in neuronal dysfunction and neurodegeneration (32). In age-matched controls, pSer129 staining was undetectable. In contrast, PD sections showed an accumulation of aSyn in the SN, with a granular pattern **(Fig. 4C)**. Quantification of aSyn^+^ cells showed a significant increase in PD samples compared to controls (****p < 0.0001; **Fig. 4D**). Together, these results confirm the loss of DAergic neurons and the pathological accumulation of aSyn in the SN of PD patients. Finally, we investigated the involvement of T-cells in PD by examining the presence of CD3^+^ T cells using immunofluorescence. We observed a marked presence of CD3^+^ cells in the SN of PD samples, while age-matched controls displayed minimal signal (**Fig. 4E**). Quantitative analysis confirmed a significant increase in CD3^+^ T-cells in PD patients compared to controls (****p < 0.0001) (**Fig. 4F**), indicating enhanced immune cell infiltration in the diseased tissue. To determine whether T-cell infiltration was associated with DAergic neuronal loss, we performed correlation analyses. CD3^+^ T-cell density was strongly and inversely correlated with the number of TH^+^ neurons (R = – 0.82, p = 6.2e-15; **Fig. 4G**), suggesting that T-cell infiltration may contribute to dopaminergic neurodegeneration. Moreover, the CD3^+^ cell count also correlated positively with aSyn levels (R = 0.86, p < 2.2e-16) **(Fig. 4H)**, consistent with previous reports that aSyn aggregates could promote chemokine release, leading to T-cell recruitment (33). Thus, our results contribute to the growing evidence that CD3^+^ T-cell infiltration is associated with both DAergic neuron loss and aSyn pathology, supporting a role for immune involvement in the pathogenesis of these core features of PD.

**Fig. 4:**
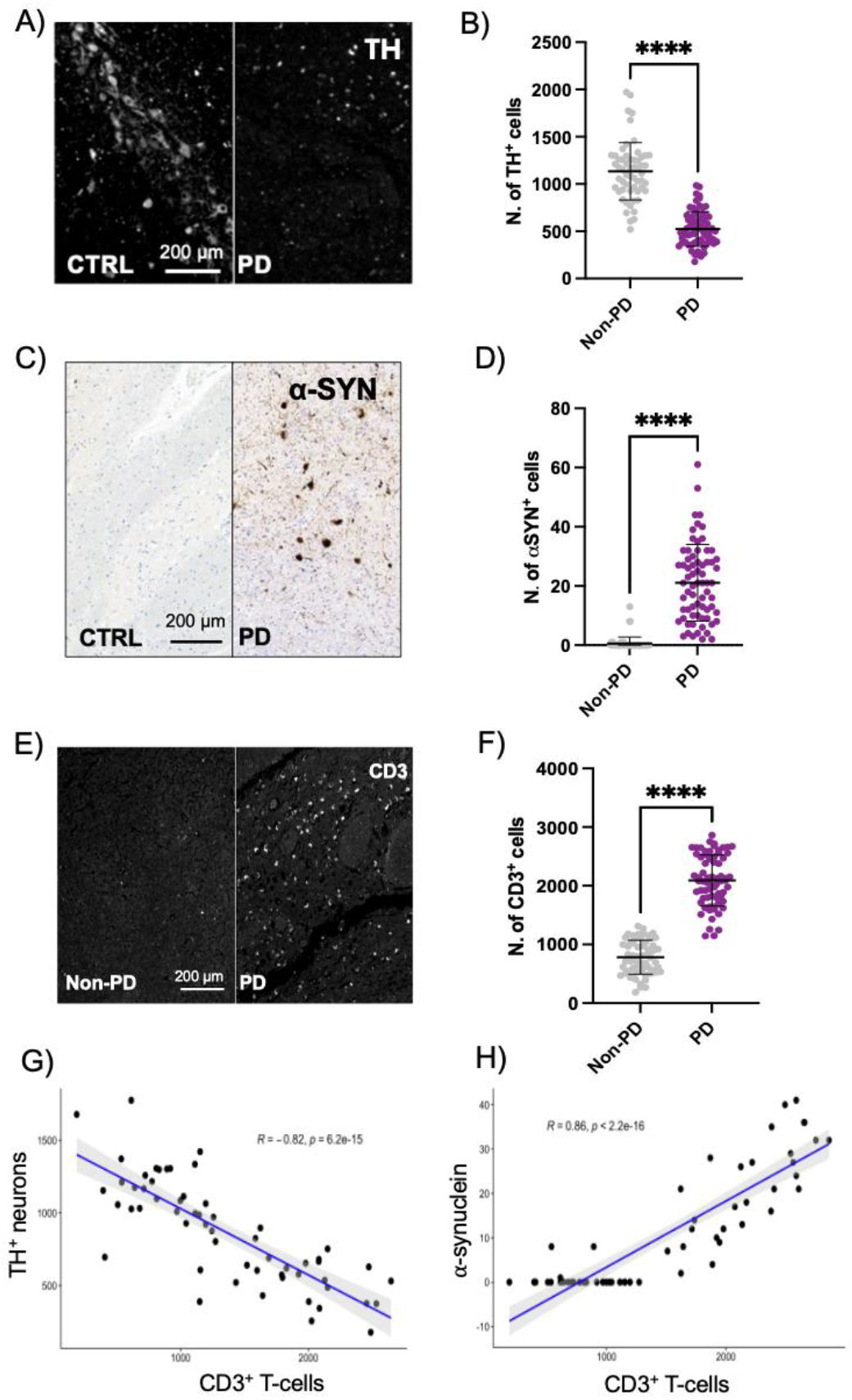
Loss of DAergic neurons, aSyn accumulation, and CD3^+^ T-cell infiltration in the substantia nigra of PD patients. **A)** Representative immunofluorescence images of SN from age-matched controls and PD patients stained for DAergic neurons (TH^+^, white); **B)** Quantification of TH^+^ neurons confirms a significant decrease in PD samples compared to age-matched controls; **C)** Representative immunohistochemical staining for aSyn from controls and PD patients; **D)** aSyn^+^ cells significantly increased in PD patients compared to the age-matched controls; **E)** Representative immunofluorescence images showing CD3^+^ T-cell infiltration in the SN of age-matched controls and PD patients; **F)** Quantification of CD3^+^ T cells that increase in PD tissues compared to the controls; **G)** Correlation analysis between CD3^+^ T-cell infiltration and the number of TH^+^ neurons; **H)** or aSyn accumulation. Statistical significance is shown in the graphs.

## DISCUSSIONS

PD is characterized by the progressive loss of DAergic neurons in the ventral tier of the SN pars compacta in the midbrain, and by intraneuronal fibrillar aSyn aggregates in Lewy bodies and neurites (34). In addition to neuronal loss and the Lewy pathology, PD shows prominent gliosis (34). Only ∼7 % of the 35,502 PD-related abstracts that we systematically parsed report a multivariate link between glial reactivity (GFAP), adaptive T-cell subsets (CD4 and CD8) and DAergic neuron loss, revealing a critical gap that our cross-species analysis fills (35–37). We therefore compared DAergic degeneration and T-cell infiltration across zebrafish, mouse models and human samples, with a particular focus on the brain and retina. Our findings revealed species- and model-specific differences in regenerative potential, glial reactivity, and T-cell infiltration, providing novel insights into the immunopathological mechanisms relevant to PD.

However, this study has limitations. Despite the value of complementary models such as MPTP-treated mice, MPTP-treated zebrafish, and L61 transgenic mice, none of these models alone fully captures the multifaceted complexity, heterogeneous pathology, and chronic progression characteristic of human PD. Additionally, the substantial species-specific differences in immune system organization and neurodegenerative capacity underscore the importance of careful interpretation (38). For example, the zebrafish’s robust ability to regenerate DAergic neurons contrasts with the largely irreversible neurodegeneration observed in mammals, potentially influencing both neuroinflammatory dynamics and tissue repair mechanisms (39). Yet, this very difference provides a valuable counterpoint: the zebrafish offer a “ceiling condition” in which injury is naturally resolved. In zebrafish, immune-glial responses are transient and pro-resolution, marked by short-lived CD4^+^ infiltration and resolving astroglial activation. In mammalian models, by contrast, inflammation is persistent and self-amplifying. This difference provides a framework to identify immune programs that enable neuronal recovery and to test whether similar pathways can be reactivated in mammals and, ultimately, in human Parkinson’s disease, a possibility that warrants further investigation. Furthermore, because human material was limited to post-mortem substantia nigra, our analysis captures only a late-stage, static view of the disease. This precludes assessment of early or evolving immune responses, which differ substantially from those observed in animal models (40). These limitations, common to preclinical research on PD, should be considered when interpreting our results.

In zebrafish, MPTP exposure caused transient SN neurodegeneration and retinal damage, followed by significant recovery. Notably, TH^+^ neuron recovery in the brain and reconstitution of the INL in the retina highlight the regenerative capacity of zebrafish, consistent with a prior study demonstrating neurogenesis in this species (13). Glial reactivity (GFAP^+^ cells) increased during the neurodegenerative phase and resolved as neuronal recovery progressed. These findings support that early astrogliosis is neuroprotective, fostering neurogenesis and structural repair as previously shown (41), and suggest that it can aid rather than hinder neuronal recovery. In contrast, αSyn-driven astroglial dysfunction may promote dopaminergic loss (42). CD3^+^ T cell infiltration mirrored this timeline, peaking at 7 dpi and declining by 14 dpi. Most were CD4^+^, whose numbers negatively correlated with TH^+^ neurons, suggesting a role in early degeneration followed by resolution that enables repair. CD8^+^ T cells were absent. Although CD4^+^ T cells are not directly cytotoxic, like CD8^+^ T cells, previous studies have shown that they can indirectly promote neuronal injury by releasing pro-inflammatory cytokines or by activating resident glial cells into a neurotoxic phenotype (43). Moreover, their decline coinciding with neuronal recovery is consistent with a regenerative role, in line with evidence that regulatory T cells can foster tissue repair (36).

MPTP-treated mice, in contrast, exhibited a DAergic degeneration accompanied by a significant increase of GFAP astrocytes without neuronal recovery. The retinal structure showed only limited reorganization, consistent with the poor regenerative capacity of the adult mammalian CNS (44, 45), where Müller glial cells and astrocytes, principal glial cells responding to injury, mount a reactive gliosis but rarely re-enter a regenerative program (45). Adaptive immunity also contributes. CD4^+^ T cells, in particular, have been implicated in aSyn-mediated neurotoxicity. Williams et al. demonstrated that CD4^+^, but not CD8^+^, T-cell depletion reduced aSyn-induced neuronal loss in PD mice, suggesting a key pathogenic role for CD4^+^ T cells infiltrating the SN (41, 43–47). Our results showed that CD8^+^ T-cell numbers negatively correlated with DAergic neuron survival, consistent with reports that MPTP induces CD8^+^ T-cell infiltration and activation (48).

The L61 transgenic model exhibits a slower, aSyn-driven loss of DAergic neurons, accompanied by astrogliosis, similar to that seen in PD patients (14). Our results demonstrated that T-cell infiltration was predominantly composed of CD4^+^ T-cells, with no detectable CD8^+^ population and CD4^+^ T cells did not correlate with neuronal loss, suggesting a modulatory rather than cytotoxic role. Similar neuroprotective functions of CD4^+^ T cells have been described in other chronic neurodegenerative conditions, such as amyotrophic lateral sclerosis (49).

Integrating these findings, our data delineate a trajectory from a transient, regeneration-permissive CD4^+^ response in zebrafish to the persistent GFAP^+^ astrogliosis and cytotoxic CD8^+^ recruitment observed in mouse models (44, 48).

Human SN analyses confirmed the cardinal PD features of DAergic neurodegeneration and aSyn aggregation (32, 34). CD3^+^ T-cell infiltration correlated positively with aSyn pathology and inversely with TH^+^ neurons counts, linking T-cell presence to both major PD hallmarks. Although we did not distinguish T-cell subsets, these findings are consistent with reports that misfolded aSyn can activate microglia and drive chemokine release, fostering recruitment of T-cells, including cytotoxic CD8^+^ populations that have been shown to damage DAergic neurons in PD (37, 50–57).

In summary, our cross-species analysis shows that transient, pro-regenerative astroglial activation in zebrafish contrasts with the persistent inflammation and dopaminergic loss seen in mammalian models and human PD. Together with published evidence for T cell-glia interactions (37, 47, 50), these findings highlight immune-glial crosstalk as a potential target for future immunomodulatory strategies.

## CONCLUSIONS

Our data reveal model-specific differences in neuronal loss, astroglial reactivity, and T cell infiltration. While zebrafish exhibited a DAergic neurodegeneration followed by regeneration, mouse models and human tissues showed DAergic neurodegeneration with no evidence of recovery.

Across 35,502 PD abstracts, <10 % connect GFAP^+^ astrogliosis, CD4^+^ or CD8^+^ T-cell dynamics and DAergic loss (35, 36). Our results help address this gap, showing that zebrafish mount a brief, CD4-dominated response after MPTP-induced degeneration, concomitant with a transient reactive gliosis (13, 41). In contrast, mammalian models and human SN display persistent GFAP^+^ reactivity and persistent T-cell presence (46, 48, 50). These findings provide a mechanistic rationale for temporally targeted immunomodulation that curbs chronic CD8^+^ and reactive gliosis while preserving early CD4 trophic support, a strategy also supported by recent murine and human evidence (37, 47).

Our results showed the dual nature of immune responses in neurodegeneration, with distinct temporal dynamics and functional outcomes depending on the model. A deeper understanding of the interplay between neurons, glial cells, and T-cells across species could provide insight into the pathophysiology of PD and open avenues for the development of targeted immunomodulatory therapies.

## Acknowledgements

The authors gratefully acknowledge Hans Ruedi Widmer for granting access to the archive of human midbrain paraffin-embedded tissues and for providing the unique opportunity to perform histological analyses with these precious samples.

