## Supplementary data for "Comparative analysis of T-cell signatures and astroglial reactivity in Parkinson’s pathology across animal models with distinct regenerative capacities"

### SUPPLEMENTARY MATERIAL

#### T-Cell involvement in DA neurodegeneration in Zebrafish

To assess T-cell infiltration following MPTP-induced injury, immunofluorescence for $CD3^+$ T cells was performed in the PT of healthy control and MPTP-treated zebrafish. Results revealed a significant increase in the number of $CD3^+$ T cells at 7 dpi compared to both control and 1 dpi (**Suppl. Fig. 1a, b),** suggesting that $CD3^+$ T cell infiltration is associated with DA neuron loss. Interestingly, at 14 dpi, $CD3^+$ T-cell numbers significantly decreased compared to 7 dpi, reaching the baseline, suggesting that T cell recruitment is linked to the acute phase of neurodegeneration and may resolve as the tissue begins to recover. A significant negative correlation **(Suppl. Fig. 1c)** was observed between the number of $CD3^+$ T cells and DA neurons ($R = –0.84$, $p < 0.0001$). This negative relationship suggests that higher levels of $CD3^+$T-cell infiltration are associated with a reduction in DA neurons, supporting the hypothesis that T cells might play an active role in promoting neurodegeneration and neuronal loss in zebrafish.

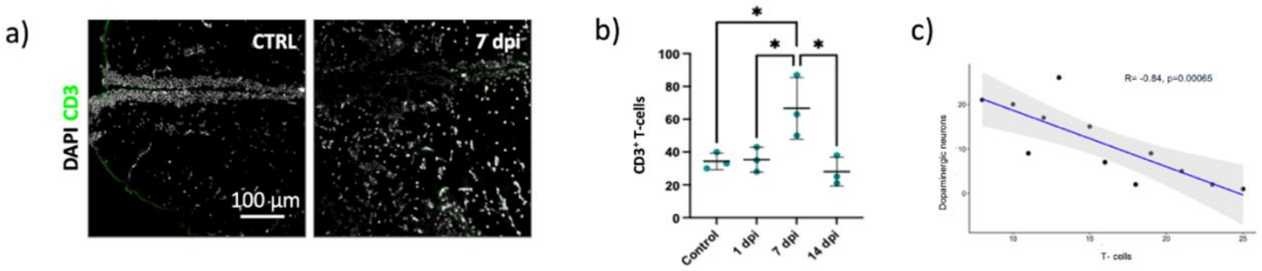


**Suppl. Fig. 1: $CD3^+$ T cells infiltration during MPTP-induced neurodegeneration. a)** Representative immunofluorescence images of the posterior tuberculum showing $CD3^+$ T cells (green) in control and MPTP-treated zebrafish at 7 dpi. Nuclei are stained with DAPI (white). **b)** Quantification of $CD3^+$ T cells at different time points after treatment. **c)** Correlation between DA neurons and the number of T-cells recruited in the MPTP-treated zebrafish brain. Statistical differences are shown in the graphs.

#### T cells immune response in L61 and MPTP treated mice

To investigate T-cell infiltration in different models of PD, we quantified $CD3^+$ T cells in the PT of L61 transgenic mice and MPTP-treated mice. In L61 mice, we observed a significant increase in $CD3^+$ T cells compared to wild-type controls **(Suppl. Fig. 2a)**, indicating T-cell recruitment in response to chronic α-synuclein accumulation. Similarly, in the MPTP model, $CD3^+$ T-cell infiltration into the PT was observed to follow a time-dependent pattern. At 1 dpi, a slight but not statistically significant increase in $CD3^+$ cells was detected compared to controls. This infiltration increased at 7 dpi and reached the maximum level at 14 dpi **(Suppl. Fig. 2b**), indicating a gradual accumulation of T cells over time. These results suggest that the adaptive immune response becomes stronger as neurodegeneration advances, with the highest level of T-cell infiltration occurring during the later stages of the neuronal damage. To explore whether T-cell presence was associated with DA neuron loss, we performed correlation analyses between $CD3^{+}$ T-cells and the number of $TH^{+}$ neurons. In L61 mice **(Suppl. Fig. 2c)**, no correlation was observed between T-cell presence and DA neuronal loss. Conversely, in the MPTP model **(Suppl. Fig. 2c)**, we observed a significant negative correlation ($R = -0.64$, $p = 0.0025$), indicating that higher numbers of $CD3^{+}$ T cells were associated with a stronger reduction in $TH^{+}$ neurons **(Suppl. Fig. 2d)**. These findings support the hypothesis that $CD3^{+}$ T cells may contribute to neurodegeneration in the MPTP model, while their role in the L61 model appears to be more complex and not linked to neuronal loss.

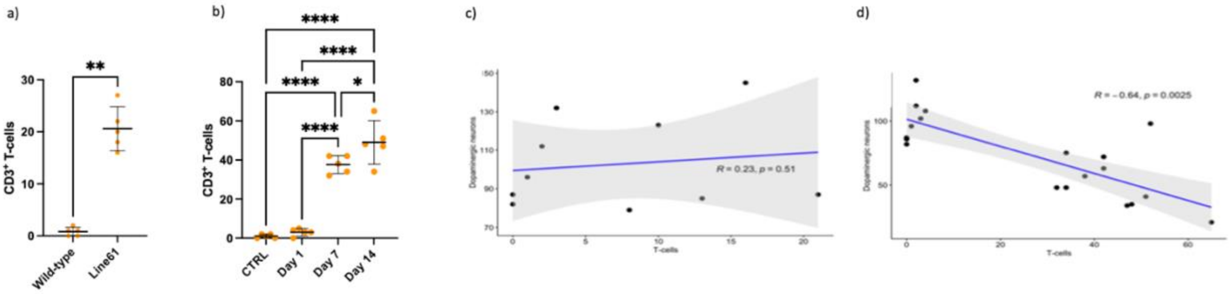


**Suppl. Fig 2: $CD3^{+}$ T-cell infiltration and correlation with DA neuron loss in L61 and MPTP models. a)** Quantification of $CD3^{+}$ T cells in the PT of L61 and **b)** MPTP-treated mice. **c)** Correlation between $CD3^{+}$ T cells and $TH^{+}$ neurons in L61 and **d)** MPTP-treated mice. Statistical differences are shown in the graphs.

**Bibliographic data mining**

**Supplementary table 1. PubMed interrogation strategy**

| Model | PubMed search strategy* |
|---|---|
| Zebrafish | (Parkinson Disease[MeSH] OR parkinson*[tiab]) AND (zebrafish[MeSH] OR zebrafish[tiab] OR Danio rerio[tiab]) AND (MPTP[tiab] OR "dopaminergic neuron"[tiab] OR retina[tiab] OR regeneration[tiab] OR "T cell"[tiab] OR astrocyte[tiab] OR microglia[tiab] OR immun*[tiab]) AND ("1960/01/01"[PDAT] : "2025/12/31"[PDAT]) |
| Mouse | (Parkinson Disease[MeSH] OR parkinson*[tiab]) AND (mouse[MeSH] OR mice[tiab] OR murine[tiab]) AND (MPTP[tiab] OR "alpha-synuclein"[tiab] OR L61[tiab] OR Thy1[tiab] OR transgenic[tiab]) AND ("T cell"[tiab] OR lymphocyte[tiab] OR immun*[tiab] OR microglia[tiab] OR astrocyte[tiab] OR retina[tiab]) AND ("1960/01/01"[PDAT] : "2025/12/31"[PDAT]) |

| | |
|---|---|
| Human | (Parkinson Disease[MeSH] OR parkinson*[tiab]) AND (human[MeSH] OR human[tiab] OR patient[tiab]) AND ("post-mortem"[tiab] OR "substantia nigra"[tiab] OR immunohistochemistry[tiab] OR "T cell"[tiab] OR lymphocyte[tiab] OR immun*[tiab] OR microglia[tiab] OR astrocyte[tiab] OR "alpha-synuclein"[tiab] OR Lewy[tiab]) AND ("1960/01/01"[PDAT] : "2025/12/31"[PDAT]) |
| Comparative | (Parkinson Disease[MeSH] OR parkinson*[tiab]) AND (zebrafish[tiab] OR Danio rerio[tiab] OR mouse[tiab] OR mice[tiab] OR murine[tiab] OR human[tiab]) AND (model*[tiab] OR comparative[tiab]) AND ("T cell"[tiab] OR lymphocyte[tiab] OR immun*[tiab] OR microglia[tiab] OR astrocyte[tiab] OR "alpha-synuclein"[tiab]) AND ("1960/01/01"[PDAT] : "2025/12/31"[PDAT]) |
